# Differential and sequential immunomodulatory role of neutrophils and Ly6C^hi^ inflammatory monocytes during antiviral antibody therapy

**DOI:** 10.1101/2020.04.22.055533

**Authors:** Jennifer Lambour, Mar Naranjo-Gomez, Myriam Boyer-Clavel, Mireia Pelegrin

## Abstract

Antiviral monoclonal antibodies (mAbs) can generate protective immunity through Fc-Fcγ Rs interactions. Using a mouse model of retroviral infection, we previously showed a crucial role for immune complexes (ICs) in the enhancement of T-cell responses through FcγR-mediated activation of dendritic cells (DCs). However, IC-FcγR interactions involve different cells of the immune system other than DCs such as neutrophils and monocytes. These two myeloid cell-types are innate effector cells rapidly recruited to sites of infection. In addition to being key cells to fight against invading pathogens, they are also endowed with immunomodulatory properties. While the role of DCs in enhancing antiviral immune responses upon mAb treatment has been addressed in several studies, the role of neutrophils and monocytes has been much less studied. Here we addressed how mAb therapy affects the functional activation of neutrophils and inflammatory monocytes in retrovirus-infected mice. We found that both cell-types activated *in vitro* by viral ICs secreted high levels of chemokines able to recruit monocytes and neutrophils themselves. Moreover, inflammatory cytokines potentiated chemokines and cytokines release by IC-activated cells and induced FcγRIV upregulation. Similarly, infection and mAb-treatment upregulated FcγRIV expression on neutrophils and inflammatory monocytes and enhanced their cytokines and chemokines secretion. Notably, upon antibody therapy neutrophils and inflammatory monocytes displayed distinct functional activation states and sequentially modulated the antiviral immune response through the secretion of Th1-type polarizing cytokines and chemokines. Our work provides novel findings on the immunomodulatory role of neutrophils and monocytes in the enhancement of immune responses upon antiviral mAb therapy.

## Introduction

The development of powerful antiviral monoclonal antibodies (mAbs) has provided new therapeutic opportunities to treat severe viral infections ^1,2^. Fc-dependent mechanisms are crucial for efficient antiviral activity of neutralizing mAbs through the engagement of IgG receptors (FcγRs) expressed on immune cells. These Fc-FcγR interactions lead to the elimination of viral particles and virus-infected cells through phagocytic and cytotoxic mechanisms (i.e. antibody-dependent cellular phagocytosis (ADCP), antibody-dependent cell-mediated cytotoxicity (ADCC),…) ^3^. Moreover, studies in different animal models of viral infection, including ours, have provided evidence that mAbs can also enhance antiviral immune responses (so called “vaccinal effects”) in a Fc-dependent manner ^4^. These vaccinal effects have been recently reported in HIV-infected patients treated with broadly neutralizing mAb (bnAbs) ^5–7^ although the mechanisms involved have not been identified thus far. The elucidation of the molecular and cellular mechanisms driving Fc-dependent, mAb-mediated immunomodulation is therefore an important issue that will be key to achieving protective immunity against severe viral infections by mAbs.

While several Fc-mediated effector functions (i.e. ADCC, ADCP, ….) have been shown to be required for antibody-mediated antiviral protection, ^8–11^ whether and how FcγR engagement by antiviral mAbs affects the immunomodulatory properties of different FcγR-expressing cells (i.e. cytokines/chemokines secretion, activation markers expression, …) has been little studied. In addition, the specific contribution of different FcγRs-expressing cells in the induction of vaccinal effects by mAbs still remains ill-understood. These issues are important to be addressed due to their potential clinical implications. However, multiple restrictions (i.e. technical and ethical issues, costs, …) largely limit those studies in humans and non-human primates (NHP). As an alternative, *in vivo* studies in immunocompetent mice infected with the Murine Leukemia Virus FrCasE allowed the identification of several immunological mechanisms that drive protective immunity upon mAb therapy ^4,12^. We have previously shown that treatment of FrCasE-infected mice with the neutralizing mAb 667 (recognizing the retroviral envelope glycoprotein, expressed on virions and infected cells) elicits protective adaptive antiviral immunity through the engagement of FcγRs ^13,14^. Notably, mAbs form immune complexes (ICs) with viral determinants that enhance antiviral T-cell responses through FcγR-mediated binding to dendritic cells (DCs) ^13,15–17^. However, IC-FcγR interactions may involve different cells of the immune system other than DCs (i.e. neutrophils, monocytes, ….). We have recently shown a key immunomodulatory role of neutrophils in the induction of protective humoral responses *via* the acquisition of B-cell helper functions (i.e. B-cell activating factor secretion) upon engagement of FcγRs by the therapeutic mAb ^18^. While the role of IC-activated DCs in the enhancement of antiviral immune responses has been addressed in several studies ^12,19,20^ the role of IC-activated neutrophils has mostly been overlooked. Neutrophils are innate effector cells rapidly recruited to sites of infection. Evidence shows that, in addition to being key effector cells to fight against invading pathogens, neutrophils are also endowed with immunomodulatory properties through the secretion of a plethora of chemokines and cytokines ^21–23^. Yet, the functional activation of neutrophils by viral ICs and the resulting effect on their immunomodulatory properties have poorly been studied in the context of antiviral mAbs therapies. Similar to neutrophils, inflammatory Ly6C^hi^ monocytes are also rapidly recruited to sites of infection and are key players to fight against viral infections ^24^. Recently, it has been reported that the therapeutic effect of antiviral mAb requires the engagement of FcγR on monocytes ^10^. This depends on Fc-FcγR interactions that are crucial to control viral spread through the enhancement of ADCP. However, the potential contribution of monocytes to the induction of vaccinal effects by antiviral mAb has not been reported. As both neutrophils and inflammatory monocytes display multiple immunomodulatory functions and can mediate protective immunity, immunopathology or immunosuppression in a context dependent manner, it is important to dissect how antiviral mAb therapy shapes the phenotype and functional properties of these FcγR-expressing cells as this might have important therapeutic implications.

Here, we used the FrCasE retroviral model to address how viral infection, with or without mAb therapy, affects the phenotypical and functional activation of neutrophils and inflammatory monocytes. Both, neutrophils and monocytes activated *in vitro* by viral determinants secreted high levels of monocyte- and neutrophil-recruiting chemokines, suggesting a self-sustaining mechanism of neutrophils and monocytes recruitment upon viral infection. *In vivo*, we have shown that viral infection and mAb-treatment shape the immunomodulatory properties of neutrophils and inflammatory monocytes. Our data show that the functional activation of both cell types differs in terms of cytokine and chemokine secretion, evolves overtime and is different in the presence or the absence of mAb-treatment. Importantly, antibody therapy leads to increased secretion of proinflammatory cytokines and chemokines that are potent inducers of Th1-biased immune responses. Our work provides hitherto unreported findings on the immunomodulatory properties of neutrophils and inflammatory monocytes in a context of antiviral mAb-therapy. These findings might help to improve mAb-based antiviral therapies by tailoring therapeutic interventions aiming at harnessing the immunomodulatory properties of these cells.

## Materials and methods

### Mice

Inbred 129/Sv/Ev mice (H-2D^b^ haplotype) were used and maintained under conventional, pathogen-free facilities at the Institut de Génétique Moléculaire de Montpellier (RAM-ZEFI). They have been used without distinction as to sex and at different ages according to experiments.

### Viral stocks

FrCasE ^25^ viral stocks were produced and stored as previously described in ^26^.

### Viral infection and immunotherapy

Eight-day-old 129/Sv/Ev mice were infected by intraperitoneal (i.p.) administration with 50 μl of a viral suspension containing 50,000 focus-forming units (FFU) and treated, or not, with 30 μg 667 mAb targeting gp70 protein of viral envelope, ^27^, 1-hour p.i. and on days 2 and 5 p.i. by i.p. administration. Mice were euthanized and spleens collected at days 8 and 14 p.i.

### Phenotypical and functional activation of FcγRIV-expressing cell from spleen ex vivo

Single-cell suspensions of splenocytes were obtained from naive, infected/non-treated and infected/treated mice at 8 and 14 days p.i. Spleen cell suspensions were obtained by mechanical dissociation in PBS, then filtered in 0,70 µm strainer. 20 % of each spleen was used for immunophenotyping by FACS, 80% left was dedicated to neutrophils and monocytes sorting. Red blood cells were lysed (ACK, Lonza) and an enrichment with biotinylated anti-B220 (BD Biosciences), anti-CD3 (BD Biosciences), following of anti-biotin Ab magnet-bead coupled (Miltenyi) and magnetic LS-columns (Miltenyi) was performed to remove spleen lymphocytes and increase the sorting efficacy. Cells were stained with specific marker of populations of interest (Ly6G-BD BioSciences, Ly6C-BioLegend, CD11b-BD BioSciences) and neutrophils (Ly6G^hi^) and inflammatory monocytes (Ly6C^hi^) were sorted (>97-98% pure) using the BD Biosciences FACSAria device. Sorted cells were cultured in 96-well plates at a density of 4×10^6^ cells/ml (1×10^6^ cells/well) in 10% FBS-containing RPMI medium for 24h, and cell-free supernatants were collected and stored at -20°C, to allow cytokines and chemokines protein release quantification.

### Flow cytometry

Organs of interest were collected to realize immunophenotyping of immune cells. Spleen cell suspensions were obtained by mechanical dissociation in PBS. BM cell suspensions were obtained by dissection and PBS-2%-FBS flushing of tibias and femurs. Cells were stained at 4°C using fluorochrome-conjugated antibodies against, CD11b (M1/70, eBioscience), CD45.2 (104, BD Biosciences), CD62L (MEL-14, BD Biosciences), CD86 (GL1, BD Biosciences), Ly6G (1A8, BD Biosciences) and Ly6C (AL-21, BioLegend). FcγRIV expression was determined using 9E9 antibody (kindly provided by Dr. Pierre Bruhns, Institut Pasteur), produced by BioXcell, and then labeled with Alexa Fluor 647. Forward scatter area and forward scatter time-of-flight, as well as side scatter, were used to remove doublets from flow cytometry analyses. Cells were analyzed on FACS LSR Fortessa (BD Bioscience), and the data were analyzed using the FlowJo software.

### Neutrophils and monocytes isolated from BM used in in vitro experiments

Neutrophils and monocytes were purified from 8 to 11-week-old naive mice BM. After dissection of lower limbs, BM cell suspensions were collected by PBS-2% FBS EDTA (2 mM) flushing (25G needle) of tibias and femurs. BM cell suspensions were filtered with a 0,40 µm nylon strainer. Two magnetic-based cell sorting (MACS) isolation kits were used to purified either neutrophils (Miltenyi Biotec) and monocytes, (Miltenyi Biotec) by negative selection, both with high purity (>97-98%), determined by FACS (LSR Fortessa, BD Bioscience). Cells were placed in culture in U bottom 96-well plates at a concentration of 1 million/ml in 10% FBS-containing RPMI medium.

### Stimulation of neutrophils and monocytes in vitro

Purified neutrophils and monocytes were seeded at 150 000 cells/ well in 150 µl of RPMI, then incubated for 24h with LPS (1µg/ml, Sigma), or FrCasE virus (MOI 5: 5 viral particles /cell), or ICs (virus FrCasE and 1µg 667 mAb), or 1µg 667 mAb alone. The MOI and 667 mAb concentration used to form ICs were previously identified by dose-response experiments involving different MOI and different 667 mAb quantities. In parallel, the same experiments were performed adding inflammatory cytokines, TNFα 100 UI/ml (Peprotech), IFNγ 100 UI/ml (eBioScience), IFNα11 a 1000 UI/ml, produced and generously provided by Dr. Gilles Uzé (DIMP, CNRS). After 24h of stimulation, supernatants were collected and stored at -20°C to quantify chemokines and cytokines protein release secretion. The viability of neutrophils and monocytes 24h post-stimulation was high (> 85%). No significant differences in cell viability were observed among the different stimulation conditions. Phenotypic activation of both neutrophils and monocytes was measured using surface markers by flow cytometry.

### Chemokines and cytokines protein release quantification

Soluble chemokines and cytokines secretion were quantified from cell-free collected supernatants of *in vitro* cultured neutrophils and monocytes and of sorted splenic neutrophils and inflammatory monocytes (of naive, infected/non-treated and infected/treated mice 8 and 14 days p.i. and immunotherapy), using bead-based immunoassays (LegendPLex, BioLegend) and analyzed on the BD Bioscience-LSR Fortessa device. The protein release quantification was established by the appropriate software (LEGENDplex™ data analysis).

### Statistic

Statistical analyses were performed using GraphPad Prism 5 (GraphPad Software). Data were expressed as mean ± SEM, and statistical significance was established using a parametric 1-way ANOVA test with Bonferroni’s multiple comparisons post-tests or non-parametric Kruskal-Wallis test with Dunn’s multiple comparisons post-test for multiple comparisons or paired Student’s *t* tests when two groups were compared. *P* values lower than 0.05 were considered as statistically significant.

### Study approval

All experimental procedures were performed in accordance with the French national animal care guidelines (CEEA-LR-12146 approval, Ethics committee of the Languedoc-Roussillon Region, Montpellier).

## Results

### Neutrophils activated by viral determinants, free or in the form of ICs, secrete high levels of chemokines able to recruit monocytes and neutrophils

We have previously shown a key role of neutrophils in the induction of long-term protective antiviral immunity upon mAb therapy of infected mice ^18^. To better characterize the phenotypical and functional activation of neutrophils by viral determinants, free or in the form of ICs, we isolated bone marrow (BM) neutrophils from naive mice and stimulated them for 24h *in vitro* with free FrCasE virions or opsonized with the 667 mAb (ICs) (Figure 1A). Free 667 mAb was used as control. Both, FrCasE virions and ICs induced a strong activation of neutrophils as shown by a higher expression of the CD11b molecule as well as an increased frequency of CD11b^hi^ CD62L^lo^ neutrophils (Figure 1B). However, IC-mediated phenotypic activation was significantly higher.

**Figure 1:**
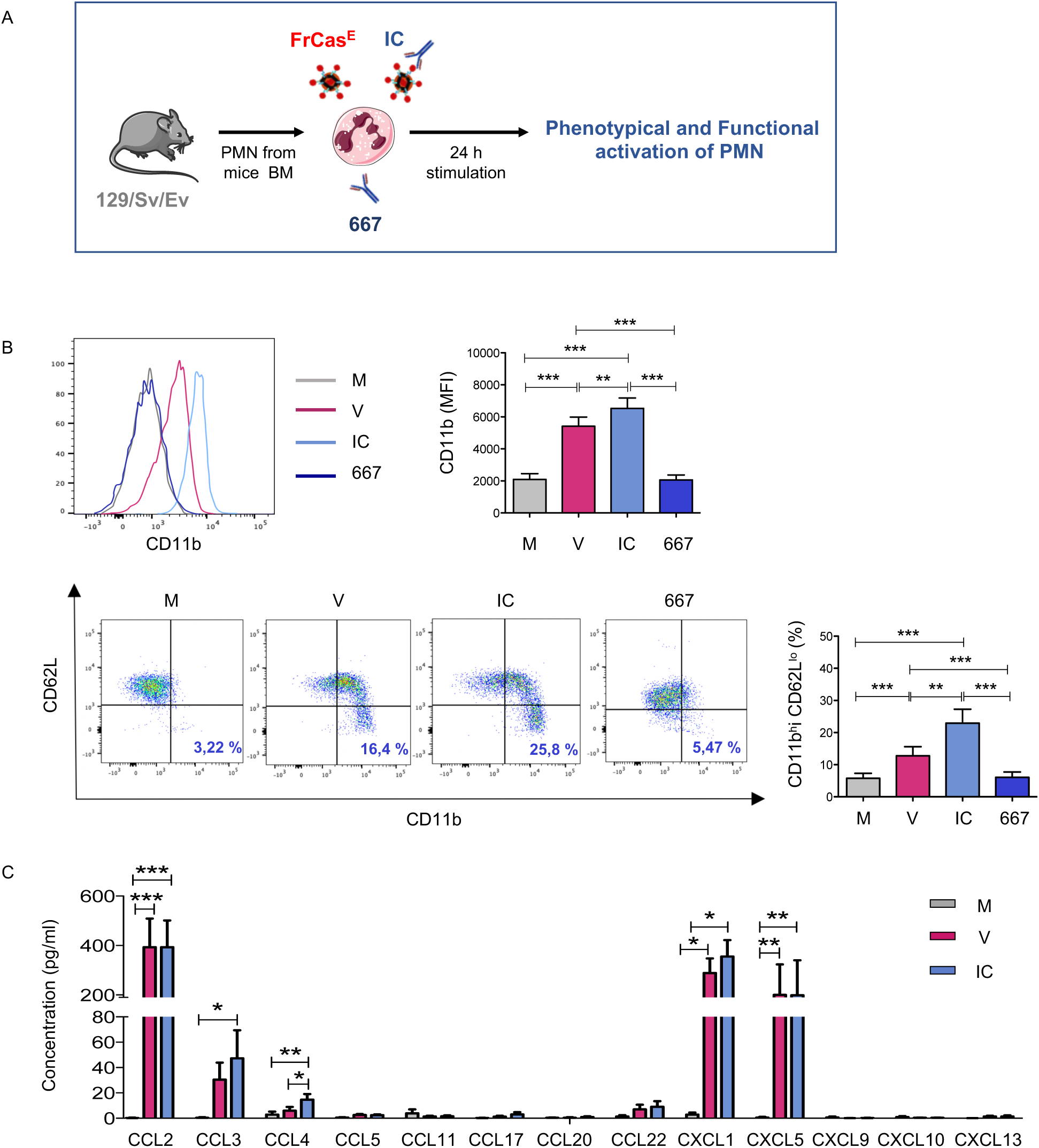
Phenotypic and functional activation of neutrophils stimulated with FrCasE virions and ICs. **A**. *Experimental scheme*. **B**. *Phenotypic activation of neutrophils stimulated by FrCasE virions or viral ICs made with the 667 mAb*. V, free virions; IC, viral ICs; M, culture medium. Free 667 mAb was used as control. Activation was assessed by monitoring CD11b expression and frequency of CD11b^hi^ CD62L^lo^ neutrophils. The data represent 12 independent experiments. **C**. *Functional activation of neutrophils stimulated by FrCasE virions (V) or viral ICs made with the 667 mAb (IC)*. Chemokines release was assessed in supernatants of neutrophils isolated from BM of naive mice (>97-98% purity) and stimulated for 24h by FrCasE virions (red) or viral ICs (blue) or left unstimulated (grey). The data represent 5 independent experiments. Data are expressed as means +/-SEM. Statistical significance was established using a parametric 1-way ANOVA test with Bonferroni’s multiple comparisons post-tests (*p < 0.05; **p < 0.01; ***p < 0.001).

We next assessed the functional activation of virus- and IC-stimulated neutrophils by measuring their capacity to secrete chemokines and cytokines. Both stimuli led to high secretion levels of several chemokines such as CCL2, CXCL1, CXCL5 and to a lesser extent CCL3 and CCL4 (Figure 1C), the secretion of the latter chemokine being significantly enhanced in IC-stimulated neutrophils. However, both the virus and ICs poorly induced the secretion of the 12 cytokines analyzed (Supplemental Figure 1A). This contrasted with the high secretion of IL-6 and TNFα pro-inflammatory cytokines observed upon lipopolysaccharide (LPS) stimulation despite similar CD11b upregulation induced by LPS and viral determinants (Supplemental Figures 1B-1C). In addition, LPS-stimulated neutrophils only secreted a low amount of CCL3 and CCL4 chemokines, with no secretion of CCL2, CXCL1, CXCL5 chemokines (Supplemental Figure 1D). These data show that bacterial-related pathogen-associated molecular patterns (PAMPs) induce a functional activation of neutrophils different from that of viral stimuli. Interestingly, the chemokines which were more strongly secreted by neutrophils upon viral stimuli (but not by LPS) have been shown to be involved in the recruitment of neutrophils themselves (CXCL1, CXCL5) and inflammatory monocytes (CCL2).

### Inflammatory monocytes activated by viral determinants, free or in the form of ICs, secrete high levels of chemokines able to recruit neutrophils and monocytes

We next assessed the activation of Ly6C^hi^ monocytes isolated from naive mice BM and stimulated for 24h with virus or ICs (Figure 2A). Both stimuli significantly activated monocytes (as depicted by an increased CD86 expression) (Figure 2B) and induced the secretion of CXCL5, CXCL1 and CCL2 chemokines, the secretion of the latter chemokine being significantly enhanced in IC-stimulated monocytes (Figure 2C). As compared to neutrophils, higher amounts of the neutrophil-recruiting chemokine CXCL1 were detected as well as lower amounts of CXCL5 and CCL2. Virus- and ICs induced a weak secretion of most of the 12 cytokines analyzed (Supplemental Figure 2A). This contrasted with high level secretion of IL-6, TNFα and IFNγ observed upon LPS stimulation (Supplemental Figure 2C). LPS stimulation also induced a wider and different panel of chemokine release (Supplemental Figure 2D), notably with the secretion of high amounts of CCL3, CCL4, CCL5, and to a lesser extent CXCL1 and CXCL10. As observed in neutrophils, these data show that viral stimuli induce a functional activation of monocytes different from that of bacterial-related PAMPs.

**Figure 2:**
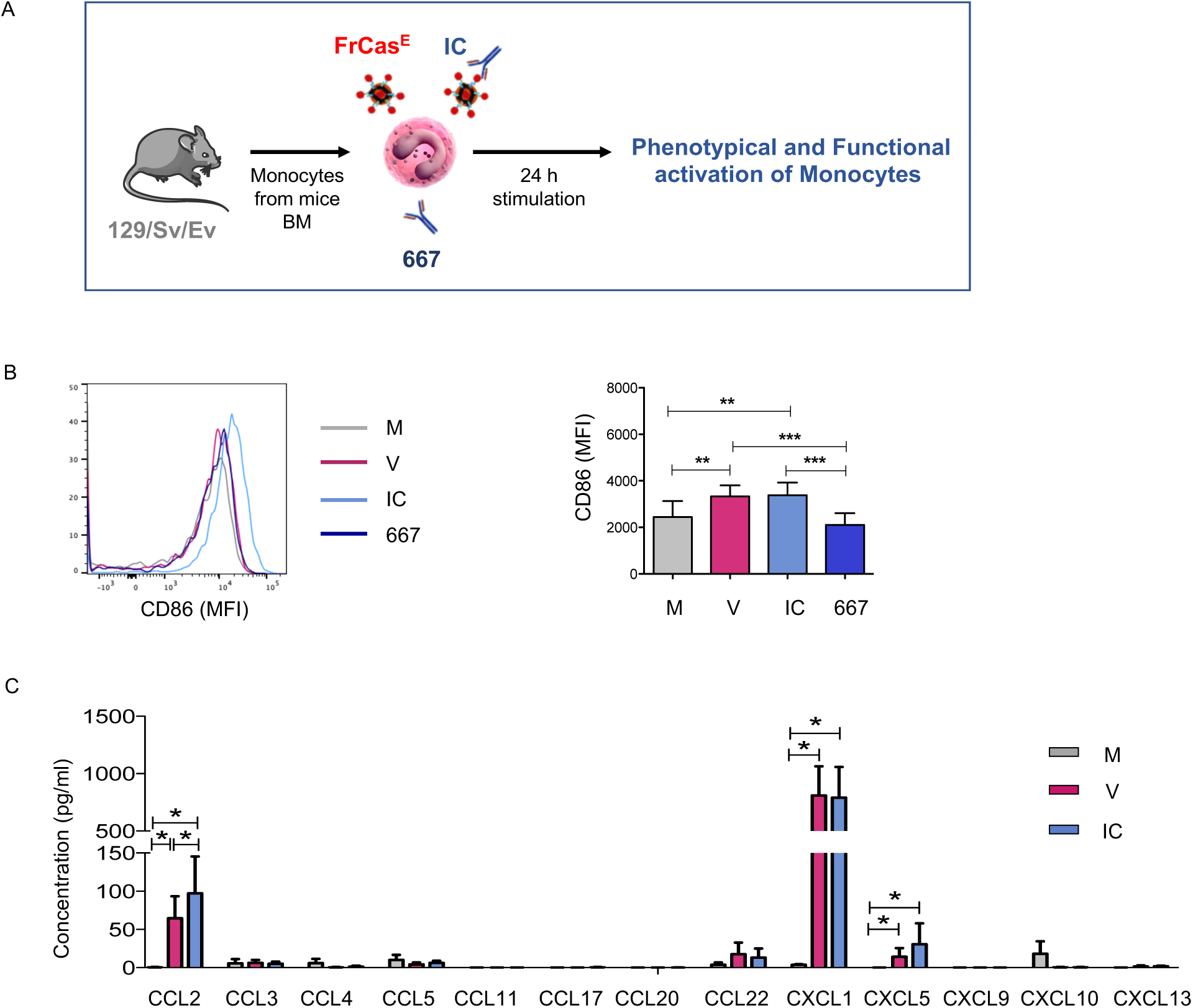
Phenotypic and functional activation of monocytes stimulated with FrCasE virions and ICs. **A**. *Experimental scheme*. **B**. *Phenotypic activation of monocytes stimulated by FrCasE virions or viral ICs made with the 667 mAb*. Free 667 mAb was used as control. V, free virions; IC, viral ICs; M, culture medium. Activation was assessed by monitoring CD86 expression. The data represent 7 independent experiments **C**. *Functional activation of monocytes stimulated by FrCasE virions (V) or viral ICs made with the 667 mAb (IC). C*hemokines release was assessed in supernatants of monocytes isolated from BM of naive mice (>97-98% purity) and stimulated for 24h by FrCasE virions (red) or viral ICs (blue) or left unstimulated (grey). The data represent 5 independent experiments. Data are expressed as means +/-SEM. Statistical significance was established using a parametric 1-way ANOVA test with Bonferroni’s multiple comparisons post-tests (*p < 0.05; **p < 0.01; ***p < 0.001).

### Inflammatory conditions potentiate the activation of neutrophils and monocytes by viral Ics

As the inflammatory microenvironment resulting from the viral infection and mAb therapy might affect the antiviral immune response, we next assessed the phenotypic and functional activation of neutrophils and monocytes by viruses and ICs under an inflammatory environment (i.e. in the presence of proinflammatory/immunomodulatory cytokines, such as TNFα, IFN-I and IFNγ).

TNFα and IFNγ significantly enhanced the phenotypic activation of neutrophils by ICs (but not by free virus) (Figure 3A). We also showed that TNFα, IFNγ and IFN-I enhanced the secretion of different cytokines and chemokines by IC-stimulated neutrophils (Figures 3B-3E). Notably, we observed an enhanced secretion of TNFα (by TNFα itself), CXCL1 (by TNFα and IFNγ), CCL4 and CXCL10 (by IFNγ and IFN-I) and CCL5 (by IFNγ). In contrast, inflammatory conditions hardly modified the secretion profile of free virus-activated neutrophils as only the secretion of the CXCL10 was induced in virus stimulated cells, consistent with the IFN-dependent induction of this chemokine (Figure 3E).

**Figure 3:**
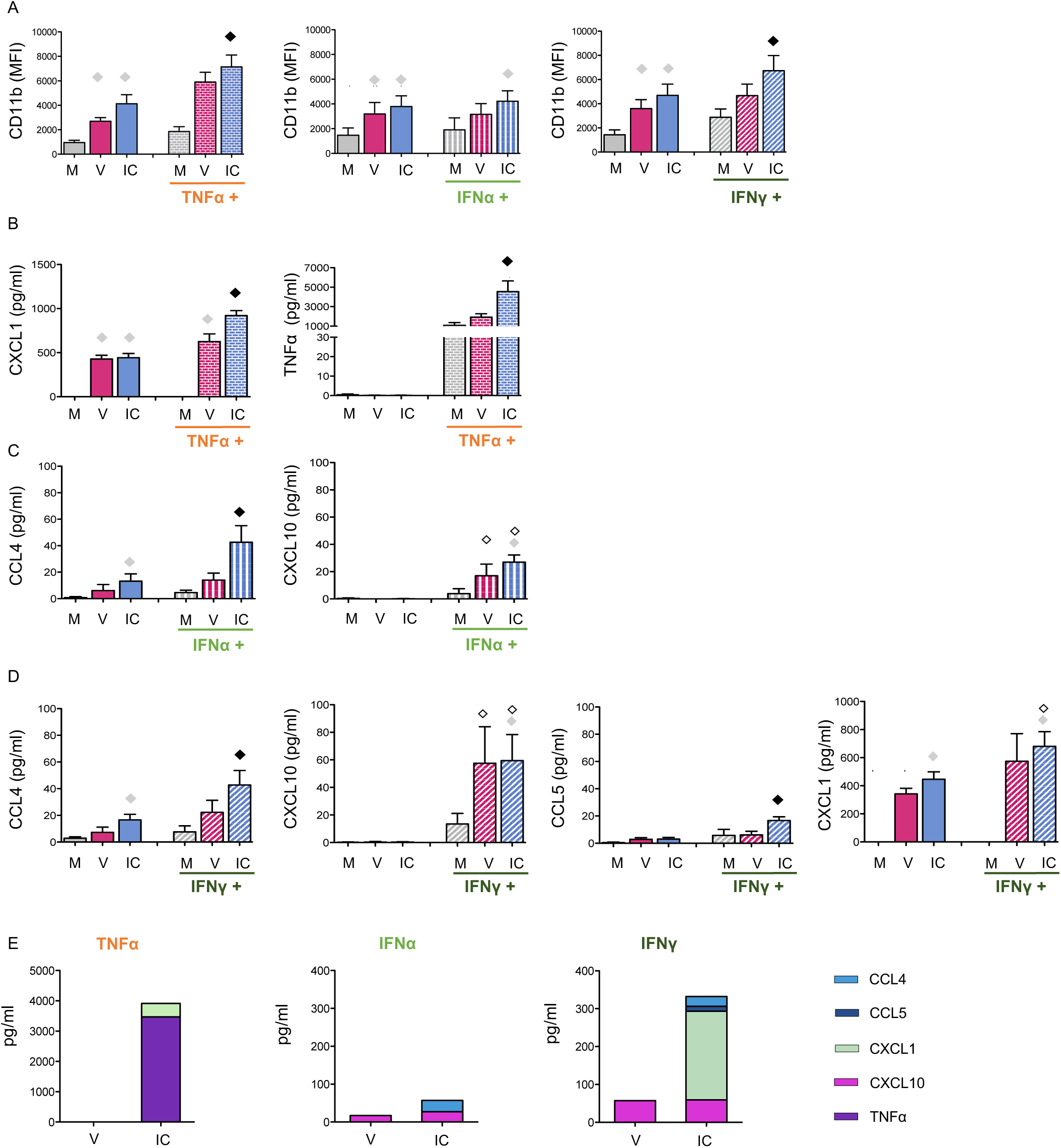
*C*ytokine stimulation potentialize the functional activation of neutrophils by ICs. BM-derived neutrophils were isolated from naive mice and activated as in **Figure 1** in the presence, or in the absence, of *TNFα, IFN-I or IFNγ*. V, free virions; IC, viral ICs; M, culture medium. **(A)**. *Phenotypic activation of activated neutrophils by TNFα, IFN-I or IFNγ*. Activation was assessed by monitoring CD11b (**A**). *Modulation of the functional activation of neutrophils by TNFα* (**B**), IFN-I (**C**) or IFNγ **(D)**. Chemokines and cytokines release were assessed in supernatants of activated neutrophils (**B**-**D**). (**E**) Histograms depict the increase in cytokine/chemokine release upon stimulation of virus- and IC-activated neutrophils by *TNFα, IFN-I or IFNγ* (calculated as the difference in the amount of chemokine/cytokine secretion by V- or IC-stimulated neutrophils in the presence or in the absence of cytokine stimulation). Only those chemokines/cytokines significantly enhanced by *TNFα* (**B**), IFN-I (**C**) or IFNγ**(D)** stimulation are depicted. The data represent 6 independent experiments for **A**, 4 independent experiments for **B-E**. Data are expressed as means +/-SEM. Diamonds indicate significant differences to all the other stimulation conditions (black diamond), or to the corresponding stimuli in the absence of cytokine stimulation (open diamond), to corresponding medium without virus or IC stimuli (grey diamond) as determined by Kruskal-Wallis test with Dunn’s multiple comparisons post-tests (p < 0.05).

With regard to monocytes, IFN-I and IFNγ significantly potentiated the phenotypic activation of IC-stimulated monocytes (as depicted by an increased expression of the CD86 molecule) (Figure 4A) but not that of virus-activated cells. Similar to neutrophils, TNFα, IFNγ and IFN-I also enhanced the secretion of different cytokines/chemokines by IC-stimulated monocytes, leading to increased secretion of CCL2 and CXCL1 (enhanced by the 3 cytokines), TNFα (enhanced by IFN-I) and CXCL10 (enhanced by both types of IFN) (Figure 4B-4E). TNFα, IFNγ and IFN-I stimulation also modulated the cytokine/chemokine secretion profile of virus-activated monocytes (i.e. CCL2, CXCL1), although to a lesser extent than IC-activated cells (Figure 4E).

**Figure 4:**
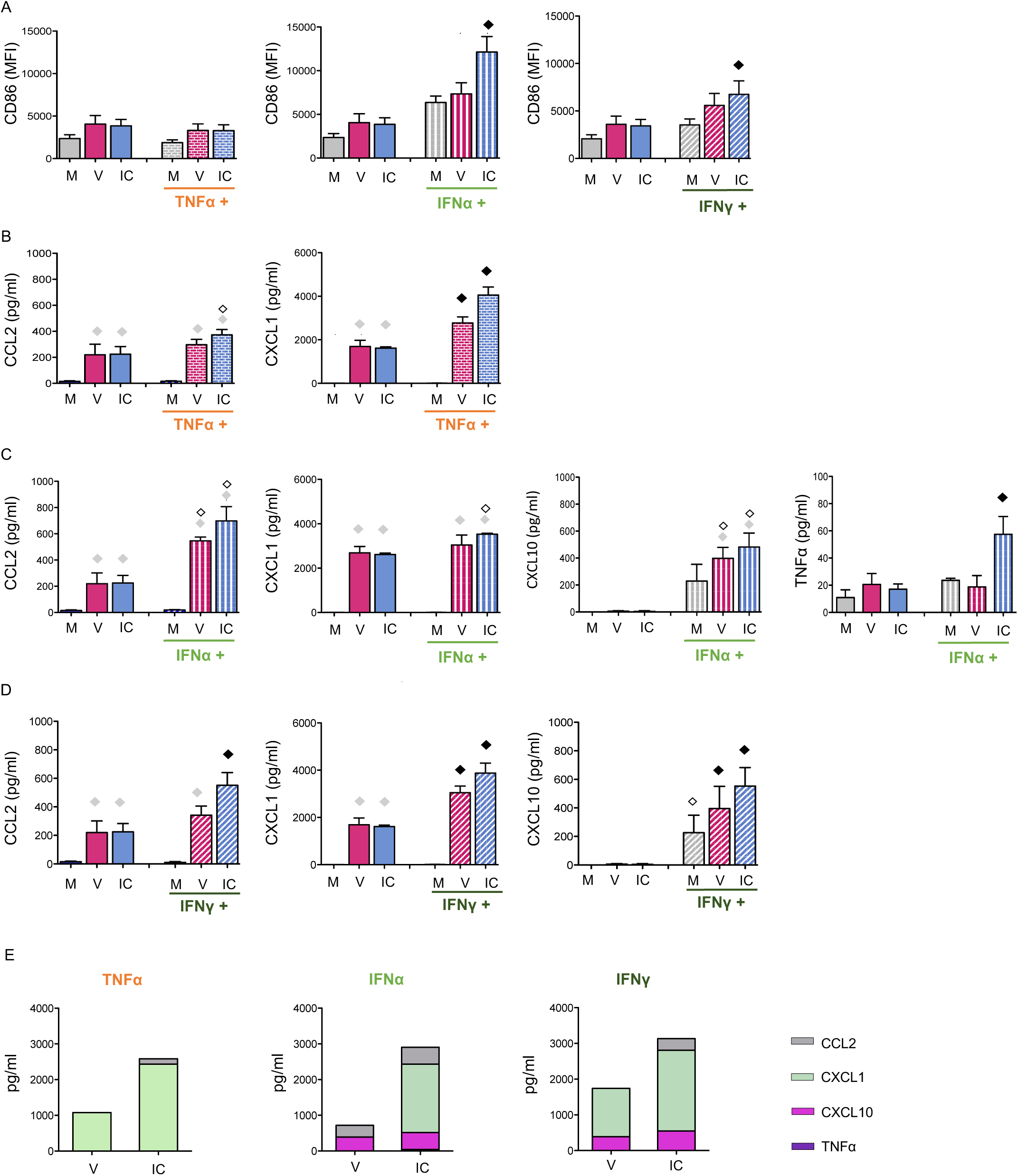
*C*ytokine stimulation potentialize the functional activation of monocytes by ICs. BM-derived monocytes were isolated from naive mice and activated as in **Figure 2** in the presence, or in the absence, of *TNFα, IFN-I or IFNγ*. V, free virions; IC, viral ICs; M, culture medium. **(A)**. *Phenotypic activation of activated monocytes (****A****) by TNFα, IFN-I or IFNγ*. Activation was assessed by monitoring CD86 expression **(A)**. *Modulation of the functional activation of monocytes by TNFα* (**B**), IFN-I (**C**) or IFNγ**(D)**. Chemokines and cytokines release were assessed in supernatants of activated neutrophils (**B**-**D**). (**E**) Histograms depict the increase in cytokine/chemokine release upon stimulation of virus- and IC-activated monocytes by *TNFα, IFN-I or IFNγ* (calculated as the difference in the amount of chemokine/cytokine secretion by V- or IC-stimulated monocytes in the presence or in the absence of cytokine stimulation). Only those chemokines/cytokines significantly enhanced by *TNFα* (**B**), IFN-I (**C**) or IFNγ**(D)** stimulation are depicted. The data represent 3 independent experiments for **A** and 3 independent experiments for **B-E**. Data are expressed as means +/-SEM. Diamonds indicate significant differences to all the other stimulation conditions (black diamond), or to the corresponding stimuli in the absence of cytokine stimulation (open diamond), or to corresponding medium without virus or IC stimuli (grey diamond) as determined by Kruskal-Wallis test with Dunn’s multiple comparisons post-tests (p < 0.05).

Altogether, these data highlight that inflammatory conditions enhance the phenotypic and functional activation of IC-stimulated neutrophils and monocytes. The cytokine/chemokine secretion enhancement observed in IC-activated cells differs depending on the stimulating cytokine and the responding cell type (i.e. increased secretion of TNFα, CCL4, CCL5 and CXCL1 by IC-activated neutrophils and TNFα, CCL2 and CXCL1 by IC-activated monocytes). It is worth to note that inflammatory conditions mainly enhanced the secretion profile of IC-activated cells but not that of virus-activated cells (notably in neutrophils).

### Inflammatory conditions upregulate the expression of FcγRIV on in vitro activated neutrophils and monocytes

FcγRs expression might differ between steady-state *versus* inflammatory/pathological conditions ^28^, however little is known about modulation of FcγRs expression in the context of viral infections and mAb-therapy. We thus assessed whether the activation of neutrophils and monocytes by virus and ICs affected the expression of the activating receptor FcγRIV, both in the absence and in the presence of inflammatory cytokines. The modulation of this FcγR is all the more relevant to be studied in this experimental model as (i) it is a high affinity receptor for IgG2a, (which is the isotype of the 667 mAb) and (ii) it is highly expressed on neutrophils, the latter having a key immunomodulatory role in mAb-mediated protection of retrovirus-infected mice ^18^. We found that IFNγ and IFN-I stimulation (but not TNF-α) led to the upregulation of FcγRIV expression on neutrophils (IFNγ) (Figure 5A) and monocytes (IFNγ and IFN-I) (Figure 5B). In contrast, in the absence of inflammatory cytokines, neither the virus nor the ICs significantly modulated the expression FcγRIV on neutrophils (Figure 5A) and monocytes (Figure 5B). However, IFNγ stimulation enhanced the upregulation of FcγRIV expression in virus- and IC-stimulated monocytes. These results show the specific effect of the different inflammatory cytokines (TNF-α, IFN-I, IFN-γ) on the modulation of FcγRIV expression.

**Figure 5:**
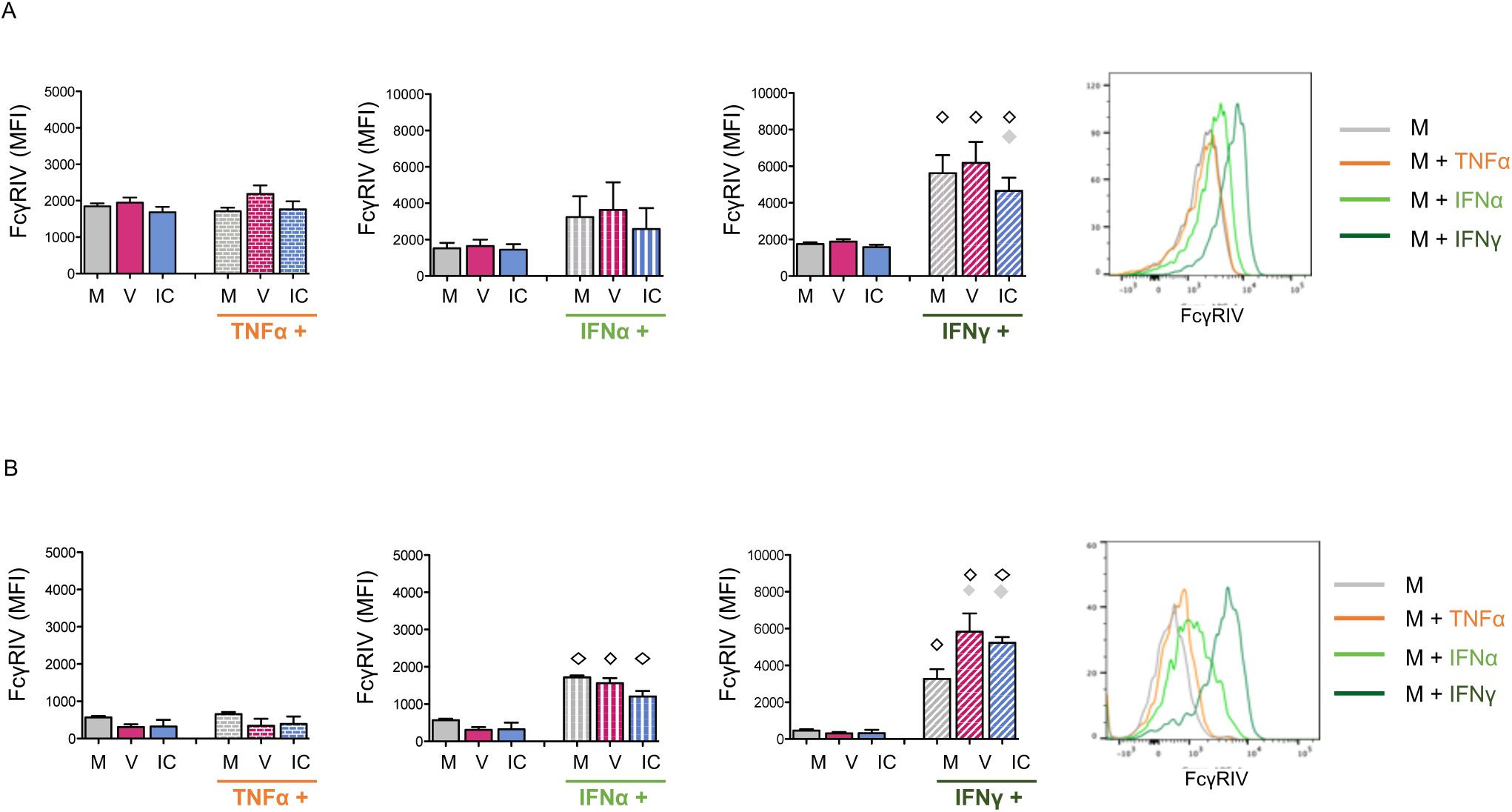
FcγRIV is upregulated by IFN stimulation on both neutrophils and monocytes cell surface. Neutrophils and monocytes were isolated from naive mice and stimulated as in **Figures 3 and 4**. FcγRIV expression was evaluated on neutrophils **(A)** and monocytes **(B)**. V, free virions; IC, viral ICs; M, culture medium. The data represent 6 independent experiments for neutrophils (**A**) and 3 independent experiments for monocytes (**B**). Data are expressed as means +/-SEM. Diamonds indicate significant differences to the corresponding stimuli in the absence of cytokine stimulation (open diamond) or to corresponding medium without virus or IC stimuli (grey diamond) as determined by Kruskal-Wallis test with Dunn’s multiple comparisons post-tests (p < 0.05).

### Viral infection and mAb therapy upregulate FcγRIV expression on neutrophils and on inflammatory monocytes

We next assessed *in vivo* whether the inflammatory environment resulting from FrCasE viral infection and 667 mAb therapy modulated the expression of FcγRIV on neutrophils and inflammatory monocytes. To this end, mice were infected and treated, or not, with the therapeutic mAb (infected/treated and infected/non-treated, respectively). Then, the expression of FcγRIV on neutrophils and inflammatory monocytes from the spleen (one of the main sites of viral replication) was evaluated at different time points post-infection (p.i.): at day 8 p.i. (i.e. corresponding to the peak of viremia) and day 14 (i.e. corresponding to the peak of primary cytotoxic T-cell responses) ^13^. Age-matched naive mice were used as controls. The cell populations of interest were defined by flow cytometry (Figure 6A) based on the expression of CD11b, Ly6G and Ly6C to gate neutrophils (CD11b^+^, Ly6G^hi^) and inflammatory monocytes (CD11b^+^Ly6G^-^Ly6C^hi^).

**Figure 6:**
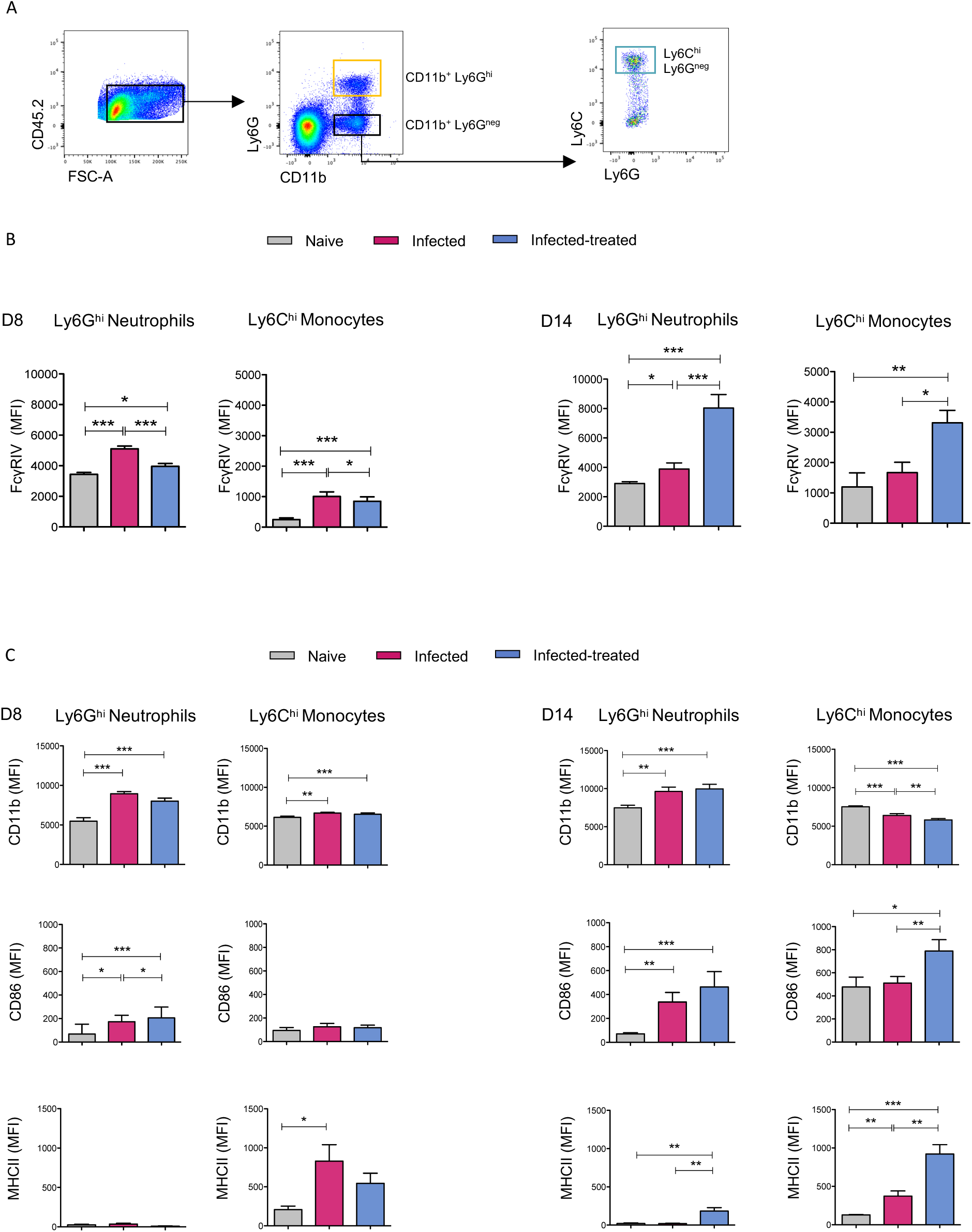
FcγRIV is upregulated *in vivo* on splenic neutrophils and on inflammatory monocytes. Splenocytes from naive (grey), infected/non-treated (I/NT; red) and infected/treated (I/T; blue) mice were analyzed on days 8 (D8) and 14 (D14) p.i. for FcγRIV expression. **(A**) Gating strategy used to define neutrophil and monocyte populations. **(B**) FcγRIV expression on *CD11b*^*+*^*Ly6G*^*hi*^ neutrophils and *Ly6C*^*hi*^ monocytes. **(C**) Cell surface markers expression (CD11b, CD86, MHCII) on *CD11b*^*+*^*Ly6G*^*hi*^ neutrophils and *CD11b*^*+*^*Ly6C*^*hi*^ monocytes surface. The data represent 5 independent experiments at D8 p.i. and 6 independent experiments at D14 p.i with at least 6-8 mice per group (I/NT and I/T) and 3-5 mice per group (naive mice). Data are expressed as means +/-SEM. Statistical significance was established using a parametric 1-way ANOVA test with Bonferroni’s multiple comparisons post-tests (*p < 0.05; **p < 0.01; ***p < 0.001).

In steady-state conditions, FcγRIV was highly expressed on splenic neutrophils. Lower FcγRIV expression was detected in Ly6C^hi^ monocytes at the different time points assessed (Figure 6B). At day 8 p.i. FcγRIV expression was significantly upregulated on neutrophils and monocytes in infected/non-treated mice. In contrast, at day 14 p.i. infected/treated mice showed a stronger upregulation of FcγRIV on both cell-types as compared to infected/non-treated and control mice.

These results suggest a distinct inflammatory environment in infected/non-treated mice *versu*s infected/treated mice that evolves over time and differentially modulates the expression of the activating FcγRIV in neutrophils and monocytes, leading to a stronger upregulation at 8 days p.i. in infected/non-treated mice and at 14 days p.i. in infected/treated mice.

### FcγRIV upregulation in infected/treated mice is associated with enhanced expression of MHC-II and co-stimulatory molecules on neutrophils and monocytes

We next addressed the activation of splenic neutrophils and inflammatory monocytes in infected mice with or without immunotherapy at days 8 and 14 p.i. by monitoring cell surface activation markers. Similar to FcγRIV expression, we showed that the activation state of the cells evolved over time. At day 8 p.i. (Figure 6C), CD11b was upregulated on neutrophils and inflammatory monocytes upon viral infection either in the absence or in the presence of immunotherapy. However, infected/treated mice showed a significantly higher CD86 upregulation in neutrophils than infected/non-treated mice. At day 14 p.i. (Figure 6C), CD11b and CD86 expression was increased in splenic neutrophils isolated from both infected/treated- and infected/non-treated mice, with no significant differences between these two groups of mice. In contrast, MHC-II upregulation was only induced in infected/treated mice, suggesting a mAb-mediated upregulation of this molecule. Similarly, we observed a significantly higher upregulation of CD86 co-stimulatory molecule and MHC-II in inflammatory monocytes from infected/treated mice as compared to infected/non-treated mice.

These data show that mAb treatment leads to the upregulation of MHC-II in neutrophils and monocytes at 14 days p.i. as well as the upregulation of the CD86 costimulatory molecule in monocytes (Figure 6C). This is associated with the FcγRIV upregulation observed in neutrophils and Ly6C^hi^ monocytes from infected/treated mice at this time point (Figure 6B).

### Neutrophils and inflammatory monocytes are differentially and sequentially activated upon antiviral mAb treatment

To further characterize the activation state of neutrophils and Ly6C^hi^ monocytes upon viral infection and mAb-therapy, we addressed the cytokine and chemokine secretion profile of splenic neutrophils and monocytes sorted from infected mice treated, or not, with the 667 mAb, at days 8 and 14 p.i. Cells sorted from age-matched naive mice were used as controls. We found that the secretion profile of neutrophils and monocytes was distinct but it was globally enhanced in both cell-types in infected/treated mice as compared to infected/non-treated mice (Figure 7A and 7B), in particular at day 8 p.i. in neutrophils and at 14 p.i. in monocytes.

**Figure 7:**
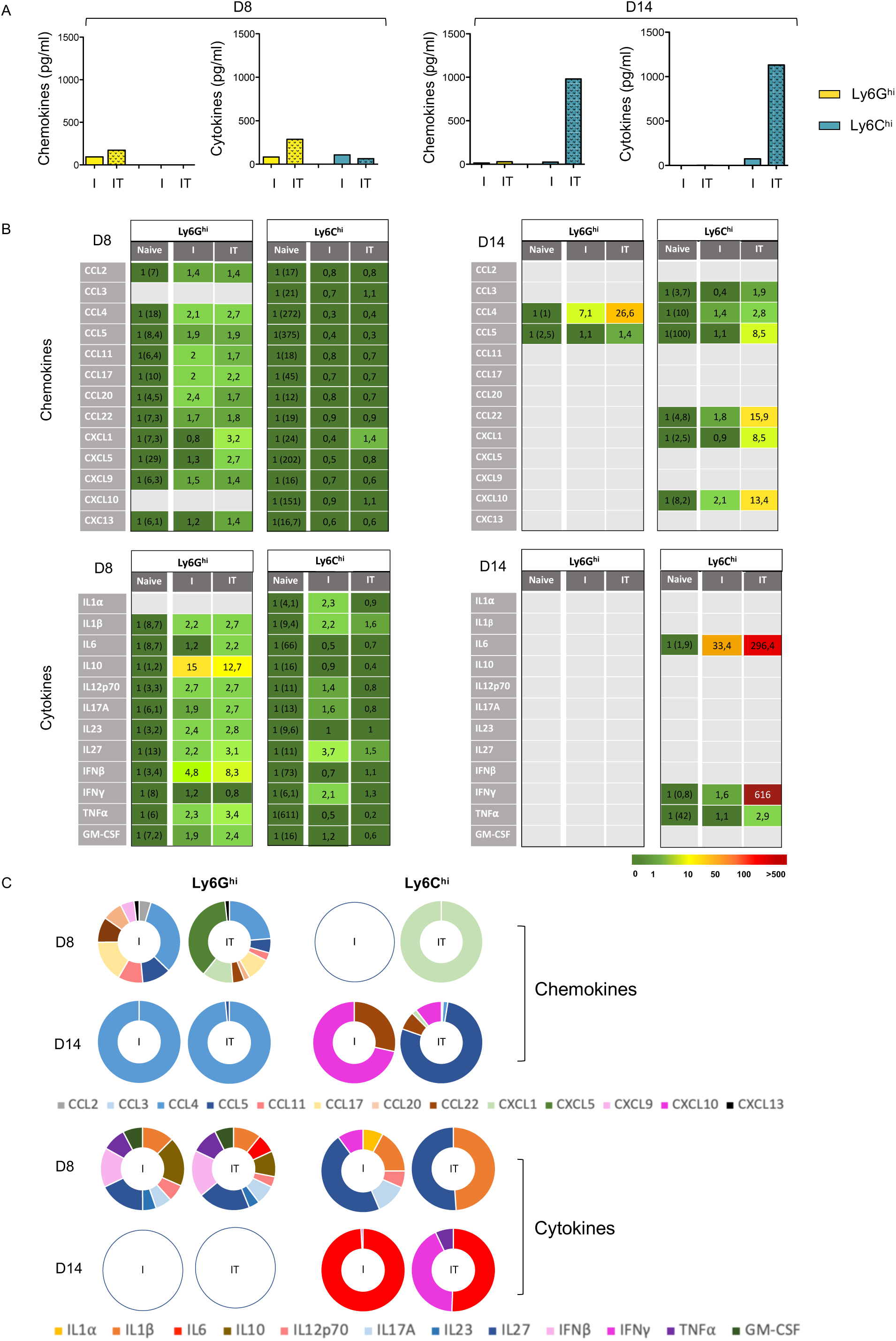
Cytokine and chemokine secretion profile of splenic neutrophils and inflammatory monocytes. Splenic neutrophils and inflammatory monocytes from naive, infected/non-treated (I) and infected/treated mice (IT) were isolated at days 8 p.i. (D8) and 14 p.i. (D14) for assaying their chemokine and cytokine secretion profile in supernatants of sorted cells cultured at a density of 4×10^6^ cells/ml (1×10^6^ cells/well) for 24 h. Data represent 3 independent experiments on D8 p.i. and 4 independent experiments on D14 p.i. with at least 6-8 mice per group. (**A**) Total quantity of chemokines and cytokines secreted by neutrophils and inflammatory monocytes from infected/non-treated (I) and infected/treated mice (I/T). (**B**) Chemokine and cytokine secretion of neutrophils and inflammatory monocytes isolated from naive, infected/non-treated (I) and infected/treated mice (IT). The color code shows fold-increase compared to secretion detected in cells isolated from naive mice (the raw chemokine and cytokine release values are given in brackets, expressed in pg/ml). (**C**) Chemokine and cytokine secretion of neutrophils and inflammatory monocytes isolated from infected/non-treated (I) and infected/treated mice (IT) expressed as percentage of chemokine and cytokine showing a fold-increase *≥*1.4 as compared to cells isolated from naive mice.

At day 8 p.i., neutrophils from both infected/non-treated and infected/treated mice showed a broad chemokine and cytokine secretion profile as deduced from an increased secretion of most of the 13 chemokines and 12 cytokines assessed, although at weak levels (Figure 7B and 7C). Notably, neutrophils from infected/treated mice showed a higher increase in IFN-I (IFN*β*) secretion (8,3-fold increase) than neutrophils from infected/non-treated mice (4,8-fold increase). In contrast, infection and mAb-treatment hardly affected the chemokine/cytokine secretion of Ly6C^hi^ monocytes as they only showed a weak increase (1.4-fold increase) in CXCL1 secretion in infected/treated mice as well as a slight increase in cytokines secretion (both in terms of diversity and fold increase) mainly in infected/non-treated mice (Figure 7B).

Interestingly, at day 14 p.i., the functional activation of both cell-types completely differed from that observed at day 8 p.i. (Figure 7B and 7C). We found a more restricted but stronger induction of chemokines/cytokines secretion, mostly in cells isolated from infected/treated mice. In contrast to the broad cytokines and chemokines secretion profile observed at day 8 p.i., neutrophils only secreted CCL4 and CCL5 chemokines and showed no cytokine secretion at day 14 p.i. It is worth noting that CCL4 secretion was strongly enhanced in infected/treated mice consistent with the enhanced secretion of this chemokine observed upon neutrophils IC-stimulation *in vitro*. In addition, Ly6C^hi^ monocytes showed a higher secretion of 6 chemokines (CCL3, CCL4, CCL5, CCL22, CXCL1 and CXCL10) and 3 Th1-polarizing cytokines (IL-6, TNFα, IFNγ) in infected/treated mice as compared to infected/non-treated mice, with notably strong induction of IL-6 and IFNγ.

In summary, FrCasE infection and 667 mAb-treatment induce the secretion of multiple chemokines and cytokines by neutrophils and monocytes. The secretion profile evolves over time in both cell-types and it is mostly increased upon antibody therapy, with notably an enhanced secretion of Th1-polarizing cytokines and chemokines by Ly6C^hi^ monocytes isolated from infected/treated mice at 14 p.i, Overall, these data suggest a role for the therapeutic mAb in the functional activation of these FcγRIV-expressing cells.

## Discussion

We have previously shown that neutrophils have a key immunomodulatory role in the induction of protective immunity by antiviral mAbs through the acquisition of B-cell helper functions ^18^. Here we provide a new insight into the immunomodulatory role of neutrophils in a context of antiviral mAb-therapy. Our work shows that antibody therapy shapes neutrophils properties, notably it leads to the upregulation of MHC-II expression and the enhanced secretion of multiple cytokines and chemokines. Our work also show that mAb-treatment strongly enhances the functional activation of inflammatory monocytes leading to the secretion of Th1-type polarizing cytokines and chemokines. These enhanced immunomodulatory functions of neutrophils and monocytes observed in infected/treated mice are associated with the upregulation of FcγRIV upon mAb treatment.

Our data show that neutrophils’ cytokine/chemokine secretion profile differs between viral *versus* bacterial stimulus. This is important to highlight as most of the studies investigating the immunomodulatory role of neutrophils have been conducted in bacterial infection models. Thus, *in vitro* stimulation of neutrophils by viral determinants led to a poor production of cytokines but to a wide and strong release of chemokine, with notably high secretion levels of the monocytes-and neutrophils-recruiting chemokines (CCL2, CXCL1, CXCL5) that was not observed upon LPS stimulation. On the contrary, LPS-stimulated neutrophils produced high amounts of proinflammatory cytokines but a narrower and weaker chemokine release. As for neutrophils, viral determinants also lead to different functional activation of monocytes than LPS. Thus, stimulation of monocytes by viral determinants (but not LPS) led to the secretion of high amounts of the neutrophils-recruiting chemokine CXCL1 and to a lesser extent the monocytes-recruiting chemokine CCL2. These results suggest a self-sustaining mechanism of neutrophils and monocytes recruitment upon viral infection, and raise the hypothesis of an early cooperation between neutrophils and monocytes in the induction of the antiviral immune response. This secretion profile of neutrophils induced by viral determinants is in agreement with an increased CCL2 release observed upon *in vitro* HIV stimulation of neutrophils ^29^. Increased levels of CCL2 and CXCL1 have also been reported in neutrophil- and monocytes-infiltrated tissues in different viral infections ^30–33^. However, chemokine increase was mostly assessed in total tissue extracts but not directly in neutrophils or monocytes isolated cells, which prevented the identification of the cell origin of chemokines. It is worth noting that a very early (i.e. 24-48h p.i.) but transient expression of CCL2 and CXCL1 has also been reported upon infection by RSV, CMV and influenza virus ^30,31,34^. Such early but transient expression would be in agreement with the very low secretion of these chemokines detected at day 8 p.i. in infected mice treated, or not, with the therapeutic mAb.

Our data also highlight the effect of inflammatory conditions on the modulation of the functional activity of neutrophils and monocytes. Thus, the *in vitro* stimulation with inflammatory/immunomodulatory cytokines potentiated the release of several chemokines and cytokines by IC-activated neutrophils and monocytes (Figure 3) (i.e. CCL4, CCL5, CXCL1, TNFα) while having a less pronounced or no effect on virus-activated cells. This is consistent with the upregulation of FcγRIV also observed in cytokine-stimulated neutrophils and monocytes. Although not formally shown, these results support a role for the inflammatory environment and IC-stimulation in the enhancement of chemokine/cytokine secretion by neutrophils and monocytes observed in infected-treated mice. Our *in vitro* observations also allowed us to dissect the specific effect of IFNγ, IFN-I and TNFα on the enhancement of the functional properties of IC-activated neutrophils and monocytes (both in terms of secretion profile and FcγRIV modulation). Notably, IFNγ and IFN-I priming (but not TNF-α), led to the upregulation of FcγRIV in neutrophils and/or monocytes, in agreement with previous reports in other experimental settings ^35–37^. This suggests a role for both types of IFN in the upregulation of FcγRIV on neutrophils and monocytes observed *in vivo* upon infection and mAb-treatment. However, FcγRIV expression was more strongly enhanced in infected/non-treated mice at day 8 p.i. and in infected/treated mice at day 14 p.i. suggesting a different and evolving inflammatory environment in both groups of mice. Similarly, the neutrophils and monocytes secretion profile was also distinct at days 8 p.i. and 14 p.i., The different activation state of these cells observed between infected/treated versus infected/non-treated mice (notably at day 14 p.i.) suggests that the control of viral propagation by the therapeutic mAb dramatically changes the inflammatory environment and the subsequent immune outcome. Thus, in addition to blunt viral propagation, Fc-mediated clearance of opsonized virus/infected cells by immune effector cells (i.e. ADCC, CDC, ADCP, …) might generate danger signals able to induce inflammation. This, together with FcγR-triggering might lead to enhanced activation of FcγR-expressing cells. Our *in vivo* results showing FcγRIV upregulation on neutrophils and monocytes are in agreement with the FcγRs modulation induced by the inflammatory conditions resulting from bacterial and IC-mediated autoimmune pathologies ^28,38^; and provide new evidence on the regulation of FcγRs expression in the specific inflammatory context of antiviral mAb-based immunotherapies, not reported thus far.

Our work provides new mechanistic insights into the hitherto underestimated role of neutrophils as key cells in the modulation of adaptive antiviral immunity upon mAb treatment. Interestingly, at steady-state conditions, neutrophils express high levels of FcγRIV. Its expression is higher than that observed in inflammatory monocytes. Thus, it is tempting to speculate that upon viral infection and mAb-treatment, the early recruitment of neutrophils, together with their high expression of FcγRIV, result in potent Fc-triggering by ICs in these cells. This might be key to initiate the modulation of immune responses through the production of multiple cytokines and chemokines able to recruit and activate multiple innate immune cells such as monocytes, NK cells, DCs and neutrophils themselves ^22^. This suggests a potential role for neutrophils as early drivers of the induction of vaccinal effects by mAbs. Supporting this hypothesis, at day 8 p.i. neutrophils showed a higher and a wider induction of chemokines and cytokines release than monocytes, while monocytes secreted strong quantities of Th-1 polarizing cytokines and chemokines at day 14 p.i. Consistent with this, a differential and sequential functional activation of myeloid cells has been recently reported by Zhang and collaborators in a model of influenza infection ^39^. By using single-cell RNA sequencing, they showed two waves of inflammation, with neutrophils being the major contributor to the first wave while macrophages generated a second wave of proinflammatory factors. However, the contribution of the different myeloid cells to the inflammation process was only assessed at the transcription level.

Our work suggests that the upregulation of FcγRIV on neutrophils and inflammatory monocytes induced by the viral infection and mAb-treatment might increase Fc-triggering by ICs leading to improved immune responses, both in terms of magnitude and quality. In agreement with this, the FcγRIV upregulation on neutrophils and monocytes in infected/treated mice observed at 14 days p.i. is associated with (i) higher expression of CD86 costimulatory molecule and MHC-II and (ii) higher secretion of cytokines and chemokines, which might enhance antiviral immune responses. The high secretion of Th1-polarizing cytokines (IL-6, TNFα, IFNγ) ^40–44^ and chemokines (CXCL10) ^45–48^ by inflammatory monocytes from infected/treated mice at day 14 p.i, argues in favor of a mAb-mediated Th1 polarization of the antiviral immune response by these cells. According to this, we previously reported that 667-mAb treatment of FrCasE infected mice leads to a Th1-biaised immune response as observed by the development of strong CD8 T-cells responses at day 14 p.i. ^13,18,49^ as well as long-lasting protective humoral immune responses predominantly of the IgG2a isotype ^13,18,26^. This Th1-type immune response observed in infected/treated mice contrasts to the non-protective immunosuppressive responses observed in infected/non-treated mice ^50^. Our results are also consistent work of Fox and collaborators showing that the therapeutic activity of mAbs against chikungunya virus (CHIK-V) requires Fc-FcγR interaction on monocytes ^10^. In agreement with our data, mAb-treatment led to higher levels of proinflammatory chemokines and cytokines in ankles of CHIK-V infected mice. However, chemokines and cytokines were assessed in total ankle samples containing multiples myeloid cell-types (including neutrophils and monocytes) but not in isolated myeloid cells. Thus, neither the effect of mAb-treatment on the secretion profile of monocytes nor the cell origin of chemokines and cytokines were addressed. Finally, it is also worth mentioning the upregulation of MHC-II molecules observed in neutrophils sorted from infected/treated mice. This observation broadens the immunomodulatory role of neutrophils upon mAb-treatment as it suggests that neutrophils might acquire antigen-presenting cell features and induce antigen-specific T-cells responses. In agreement with this, it has been shown that upon *ex vivo* stimulation with IgG immune complex (IC) or viral cognate antigens, neutrophils upregulated expression of MHCII and costimulatory molecules and increased T cell activation ^51–53^.

Our work shows a distinct secretion profile of neutrophils and Ly6C^hi^ monocytes, in terms of type, amount and kinetics of chemokines/cytokines secretion. This highlights a differentially and sequentially activation of these FcγRIV-expressing cells upon antiviral mAb treatment and suggests their key and complementary action in the induction of protective immune responses by mAb. This is important to keep into consideration as thus far, mostly DC have been considered as key cells involved in the induction of vaccinal effects by mAbs ^13,16,17^. Thus, the identification of neutrophils and inflammatory monocytes as players in the induction of protective immunity by mAbs might have therapeutic implications. These findings might help to improve mAb-based antiviral therapies by tailoring therapeutic interventions aiming at harnessing the immunomodulatory properties of these cells. To this end, different approaches could be envisaged: (i) the use of cell-type specific, appropriate immunostimulatory host-directed therapies and (ii) the design of antiviral mAbs engineered to enhance their affinity for FcγRs expressed on human neutrophils and monocytes, such as FcγRIIa and FcγRIIIa ^17,54–56^. Thus, in addition to allow superior antibody-mediated phagocytosis, these Fc-engineered mAbs could also modulate cytokine/chemokine production to ultimately lead to more effective adaptive immune responses.

## Author contributions

Jennifer Lambour (JL), Mar Naranjo-Gomez (MN-G) and Mireia Pelegrin (MP) defined the research program. JL, MN-G, and Myriam Boyer-Clavel (MB-C) performed the experiments. JL, MN-G and MP carried out the data analyses. JL and MP wrote the manuscript. Grants to MP funded the study.

## Acknowledgments

This work was supported by grants from the ANRS (France REcherche Nord&sud Sida-hiv Hépatites; ECTZ46143; ECTZ47079), the Ligue Régionale Contre le Cancer (157935, R19063FF-RAB19013FFA), Sidaction (BI25-1-02278, A014-2-AEQ-08-01) and the Fondation pour la Recherche Médicale (SPF20120523949). M. Naranjo-Gomez, J. Lambour, and M. Pelegrin are members of the “MabImprove Labex”, a public grant overseen by the French National Research Agency (ANR) as part of the “Investments for the future” program (reference: ANR-10-LABX -53-01) that also supported this work. We thank the imaging facility MRI, which is part of the UMS BioCampus Montpellier and a member of the national infrastructure France-BioImaging, supported by the French National Research Agency (ANR-10-INBS-04, “Investments for the future”). We are grateful to the animal facility of the Institut de Génétique Moléculaire de Montpellier (RAM-ZEFI) which is part of the “Réseau des Animaleries Montpelliéraines” RAMIBiSA Facility for animal experiments. We are grateful to S. Gailhac from MRI for support in cytometry experiments, to Thierry Gostan (SERENAD Complex Biological Data Analysis Service) for support in statistical analyses, to Helen Phillips Bevis for English editing services and to Drs. Valerie Dardalhon, Gilles Uzé and Pascale Plence for critical reading of the manuscript.

## References

1. Dibo M, Battocchio EC, dos Santos Souza LM, et al. Antibody Therapy for the Control of Viral Diseases: An Update. CPB. 2019;20(13):1108–1121.

2. Salazar G, Zhang N, Fu T-M, An Z. Antibody therapies for the prevention and treatment of viral infections. NPJ Vaccines. 2017;2:19.

3. Lu LL, Suscovich TJ, Fortune SM, Alter G. Beyond binding: antibody effector functions in infectious diseases. Nat Rev Immunol. 2018;18(1):46–61.

4. Pelegrin M, Naranjo-Gomez M, Piechaczyk M. Antiviral Monoclonal Antibodies: Can They Be More Than Simple Neutralizing Agents? Trends Microbiol. 2015;23(10):653–665.

5. Naranjo-Gomez M, Pelegrin M. Vaccinal effect of HIV-1 antibody therapy: Current Opinion in HIV and AIDS. 2019;14(4):325–333.

6. Niessl J, Baxter AE, Mendoza P, et al. Combination anti-HIV-1 antibody therapy is associated with increased virus-specific T cell immunity. Nat Med. 2020;26(2):222–227.

7. Schoofs T, Klein F, Braunschweig M, et al. HIV-1 therapy with monoclonal antibody 3BNC117 elicits host immune responses against HIV-1. Science. 2016;352(6288):997–1001.

8. Bournazos S, Klein F, Pietzsch J, Seaman MS, Nussenzweig MC, Ravetch JV. Broadly Neutralizing Anti-HIV-1 Antibodies Require Fc Effector Functions for In Vivo Activity. Cell. 2014;158(6):1243–1253.

9. Earnest JT, Basore K, Roy V, et al. Neutralizing antibodies against Mayaro virus require Fc effector functions for protective activity. Journal of Experimental Medicine. 2019;216(10):2282–2301.

10. Fox JM, Roy V, Gunn BM, et al. Optimal therapeutic activity of monoclonal antibodies against chikungunya virus requires Fc-FcγR interaction on monocytes. Sci Immunol. 2019;4(32):eaav5062.

11. Gunn BM, Yu W-H, Karim MM, et al. A Role for Fc Function in Therapeutic Monoclonal Antibody-Mediated Protection against Ebola Virus. Cell Host & Microbe. 2018;24(2):221–233.e5.

12. Lambour J, Naranjo-Gomez M, Piechaczyk M, Pelegrin M. Converting monoclonal antibody-based immunotherapies from passive to active: bringing immune complexes into play. Emerg Microbes Infect. 2016;5(8):e92.

13. Michaud H-A, Gomard T, Gros L, et al. A crucial role for infected-cell/antibody immune complexes in the enhancement of endogenous antiviral immunity by short passive immunotherapy. PLoS Pathog. 2010;6(6):e1000948.

14. Nasser R, Pelegrin M, Michaud H-A, Plays M, Piechaczyk M, Gros L. Long-lasting protective antiviral immunity induced by passive immunotherapies requires both neutralizing and effector functions of the administered monoclonal antibody. J Virol. 2010;84(19):10169–10181.

15. Celis E, Chang TW. HBsAg-serum protein complexes stimulate immune T lymphocytes more efficiently than do pure HBsAg. Hepatology. 1984;4(6):1116–1123.

16. Yamamoto T, Iwamoto N, Yamamoto H, et al. Polyfunctional CD4+ T-cell induction in neutralizing antibody-triggered control of simian immunodeficiency virus infection. J Virol. 2009;83(11):5514–5524.

17. Bournazos S, Corti D, Virgin HW, Ravetch JV. Fc-optimized antibodies elicit CD8 immunity to viral respiratory infection. Nature. Published online October 8, 2020.

18. Naranjo-Gomez M, Lambour J, Piechaczyk M, Pelegrin M. Neutrophils are essential for induction of vaccine-like effects by antiviral monoclonal antibody immunotherapies. JCI Insight. 2018;3(9).

19. Wang X-Y, Wang B, Wen Y-M. From therapeutic antibodies to immune complex vaccines. npj Vaccines. 2019;4(1):2.

20. Wen Y-M, Mu L, Shi Y. Immunoregulatory functions of immune complexes in vaccine and therapy. EMBO Mol Med. 2016;8(10):1120–1133.

21. Mantovani A, Cassatella MA, Costantini C, Jaillon S. Neutrophils in the activation and regulation of innate and adaptive immunity. Nat Rev Immunol. 2011;11(8):519–531.

22. Tecchio C, Cassatella MA. Neutrophil-derived chemokines on the road to immunity. Semin Immunol. 2016;28(2):119–128.

23. Tamassia N, Bianchetto-Aguilera F, Arruda-Silva F, et al. Cytokine production by human neutrophils: revisiting the “dark side of the moon.” Eur J Clin Invest. Published online May 17, 2018:e12952.

24. Stegelmeier AA, van Vloten JP, Mould RC, et al. Myeloid Cells during Viral Infections and Inflammation. Viruses. 2019;11(2):168.

25. Portis JL, Czub S, Garon CF, McAtee FJ. Neurodegenerative disease induced by the wild mouse ecotropic retrovirus is markedly accelerated by long terminal repeat and gag-pol sequences from nondefective Friend murine leukemia virus. J Virol. 1990;64(4):1648–1656.

26. Gros L, Dreja H, Fiser AL, Plays M, Pelegrin M, Piechaczyk M. Induction of long-term protective antiviral endogenous immune response by short neutralizing monoclonal antibody treatment. J Virol. 2005;79(10):6272–6280.

27. McAtee FJ, Portis JL. Monoclonal antibodies specific for wild mouse neurotropic retrovirus: detection of comparable levels of virus replication in mouse strains susceptible and resistant to paralytic disease. J Virol. 1985;56(3):1018–1022.

28. Wang Y, Jönsson F. Expression, Role, and Regulation of Neutrophil Fcγ Receptors. Front Immunol. 2019;10:1958. doi:10.3389/fimmu.2019.01958

29. Yoshida T, Kobayashi M, Li X-D, Pollard RB, Suzuki F. Inhibitory effect of glycyrrhizin on the neutrophil-dependent increase of R5 HIV replication in cultures of macrophages. Immunol Cell Biol. 2009;87(7):554–558.

30. Hokeness KL, Kuziel WA, Biron CA, Salazar-Mather TP. Monocyte Chemoattractant Protein-1 and CCR2 Interactions Are Required for IFN- / -Induced Inflammatory Responses and Antiviral Defense in Liver. The Journal of Immunology. 2005;174(3):1549–1556.

31. Miller AL, Bowlin TL, Lukacs NW. Respiratory Syncytial Virus–Induced Chemokine Production: Linking Viral Replication to Chemokine Production In Vitro and In Vivo. J INFECT DIS. 2004;189(8):1419–1430.

32. Rubio N, Sanz-Rodriguez F. Induction of the CXCL1 (KC) chemokine in mouse astrocytes by infection with the murine encephalomyelitis virus of Theiler. Virology. 2007;358(1):98–108.

33. Seo S-U, Kwon H-J, Ko H-J, et al. Type I interferon signaling regulates Ly6C(hi) monocytes and neutrophils during acute viral pneumonia in mice. PLoS Pathog. 2011;7(2):e1001304.

34. Wareing MD, Lyon AB, Lu B, Gerard C, Sarawar SR. Chemokine expression during the development and resolution of a pulmonary leukocyte response to influenza A virus infection in mice. Journal of Leukocyte Biology. 2004;76(4):886–895.

35. Dahal LN, Dou L, Hussain K, et al. STING Activation Reverses Lymphoma-Mediated Resistance to Antibody Immunotherapy. Cancer Res. 2017;77(13):3619–3631.

36. Lehmann B, Biburger M, Brückner C, et al. Tumor location determines tissue-specific recruitment of tumor-associated macrophages and antibody-dependent immunotherapy response. Sci Immunol. 2017;2(7):eaah6413.

37. Pricop L, Redecha P, Teillaud JL, et al. Differential modulation of stimulatory and inhibitory Fc gamma receptors on human monocytes by Th1 and Th2 cytokines. J Immunol. 2001;166(1):531–537.

38. Bruhns P. Properties of mouse and human IgG receptors and their contribution to disease models. Blood. 2012;119(24):5640–5649.

39. Zhang J, Liu J, Yuan Y, et al. Two waves of pro-inflammatory factors are released during the influenza A virus (IAV)-driven pulmonary immunopathogenesis. Sant AJ, ed. PLoS Pathog. 2020;16(2):e1008334.

40. Czarniecki CW. The role of tumor necrosis factor in viral disease. Antiviral Research. 1993;22(4):223–258.

41. Hildenbrand B, Lorenzen D, Sauer B, et al. IFN-y enhances T(H)1 polarisation of monocyte-derived dendritic cells matured with clinical-grade cytokines using serum-free conditions. Anticancer Res. 2008;28(3A):1467–1476.

42. Jin P, Zhao Y, Liu H, et al. Interferon-γ and Tumor Necrosis Factor-α Polarize Bone Marrow Stromal Cells Uniformly to a Th1 Phenotype. Sci Rep. 2016;6(1):26345.

43. Uciechowski P, Dempke WCM. Interleukin-6: A Masterplayer in the Cytokine Network. Oncology. 2020;98(3):131–137.

44. Wang JP, Kurt-Jones EA, Shin OS, Manchak MD, Levin MJ, Finberg RW. Varicella-Zoster Virus Activates Inflammatory Cytokines in Human Monocytes and Macrophages via Toll-Like Receptor 2. Journal of Virology. 2005;79(20):12658–12666.

45. Gaylo-Moynihan A, Prizant H, Popović M, et al. Programming of Distinct Chemokine-Dependent and -Independent Search Strategies for Th1 and Th2 Cells Optimizes Function at Inflamed Sites. Immunity. 2019;51(2):298-309.e6.

46. Kelly-Scumpia KM, Scumpia PO, Delano MJ, et al. Type I interferon signaling in hematopoietic cells is required for survival in mouse polymicrobial sepsis by regulating CXCL10. The Journal of Experimental Medicine. 2010;207(2):319–326.

47. Tannenbaum CS, Tubbs R, Armstrong D, Finke JH, Bukowski RM, Hamilton TA. The CXC chemokines IP-10 and Mig are necessary for IL-12-mediated regression of the mouse RENCA tumor. J Immunol. 1998;161(2):927–932.

48. Tokunaga R, Zhang W, Naseem M, et al. CXCL9, CXCL10, CXCL11/CXCR3 axis for immune activation – A target for novel cancer therapy. Cancer Treatment Reviews. 2018;63:40–47.

49. Gros L, Pelegrin M, Michaud H-A, et al. Endogenous cytotoxic T-cell response contributes to the long-term antiretroviral protection induced by a short period of antibody-based immunotherapy of neonatally infected mice. J Virol. 2008;82(3):1339–1349.

50. Nasser R, Pelegrin M, Plays M, Gros L, Piechaczyk M. Control of regulatory T cells is necessary for vaccine-like effects of antiviral immunotherapy by monoclonal antibodies. Blood. 2013;121(7):1102–1111.

51. Lok LSC, Dennison TW, Mahbubani KM, Saeb-Parsy K, Chilvers ER, Clatworthy MR. Phenotypically distinct neutrophils patrol uninfected human and mouse lymph nodes. Proc Natl Acad Sci USA. 2019;116(38):19083–19089.

52. Meinderts SM, Baker G, van Wijk S, et al. Neutrophils acquire antigen-presenting cell features after phagocytosis of IgG-opsonized erythrocytes. Blood Advances. 2019;3(11):1761–1773.

53. Vono M, Lin A, Norrby-Teglund A, Koup RA, Liang F, Loré K. Neutrophils acquire the capacity for antigen presentation to memory CD4+ T cells in vitro and ex vivo. Blood. 2017;129(14):1991–2001.

54. Bournazos S, DiLillo DJ, Goff AJ, Glass PJ, Ravetch JV. Differential requirements for FcγR engagement by protective antibodies against Ebola virus. Proc Natl Acad Sci USA. 2019;116(40):20054–20062.

55. Chen TF, Sazinsky SL, Houde D, et al. Engineering Aglycosylated IgG Variants with Wild-Type or Improved Binding Affinity to Human Fc Gamma RIIA and Fc Gamma RIIIAs. Journal of Molecular Biology. 2017;429(16):2528–2541.

56. DiLillo DJ, Ravetch JV. Differential Fc-Receptor Engagement Drives an Anti-tumor Vaccinal Effect. Cell. 2015;161(5):1035–1045.

