## Supplementary material for "Differential and sequential immunomodulatory role of neutrophils and Ly6C^hi^ inflammatory monocytes during antiviral antibody therapy": Supplental figures 1 and 2

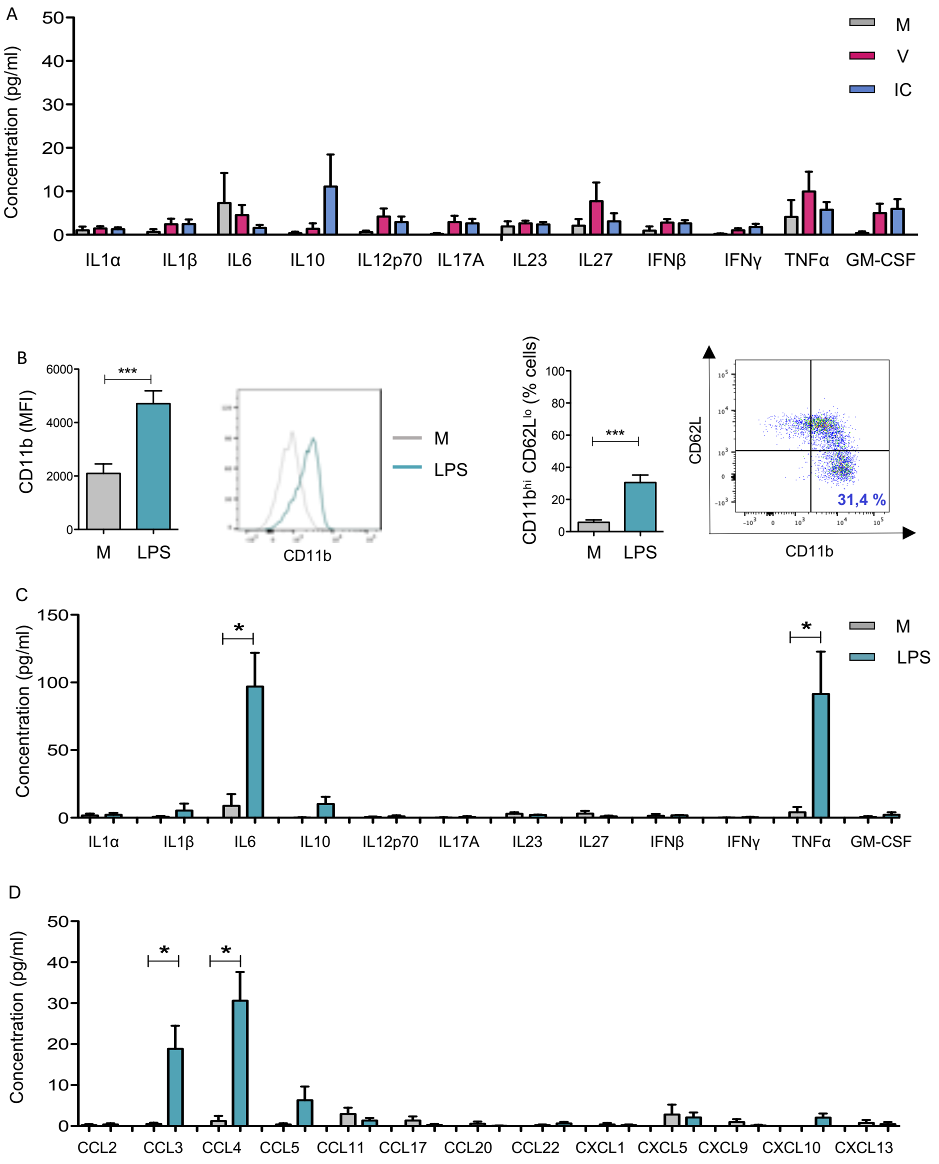


***Supplemental Figure 1.*** **Functional activation of neutrophils stimulated with viral determinants or LPS.** BM-derived neutrophils were isolated from naive mice and stimulated for 24h *in vitro* with viral determinants (virus or ICs) or LPS (1$\mu$g/ml). V, free virions; IC, viral ICs; M, culture medium **A**. *Cytokine secretion profile of neutrophils stimulated by virus (red) o viral ICs (blue) or unstimulated (grey)*. Cytokine release was assessed in supernatants of neutrophils isolated from BM of naive mice (>97-98% purity). Data represent 5 independent experiments and are expressed as means +/- SEM. Statistical significance was established using a parametric 1-way ANOVA test with Bonferroni's multiple comparisons post-tests (*p < 0.05; **p < 0.01; ***p < 0.001). **B-D**. *Functional activation of neutrophils stimulated by LPS*. Activation was assessed by monitoring CD11b expression and frequency of CD11b$^{hi}$ CD62L$^{lo}$ neutrophils (**B**) as well as the cytokines (**C**) and chemokines (**D**) released in supernatants of neutrophils stimulated for 24 h by LPS (green) or left unstimulated (grey). Data represent 12 independent experiments and are expressed as means +/- SEM. Statistical significance was established using a paired Student's t test (*p < 0.05; **p < 0.01; ***p < 0.001).

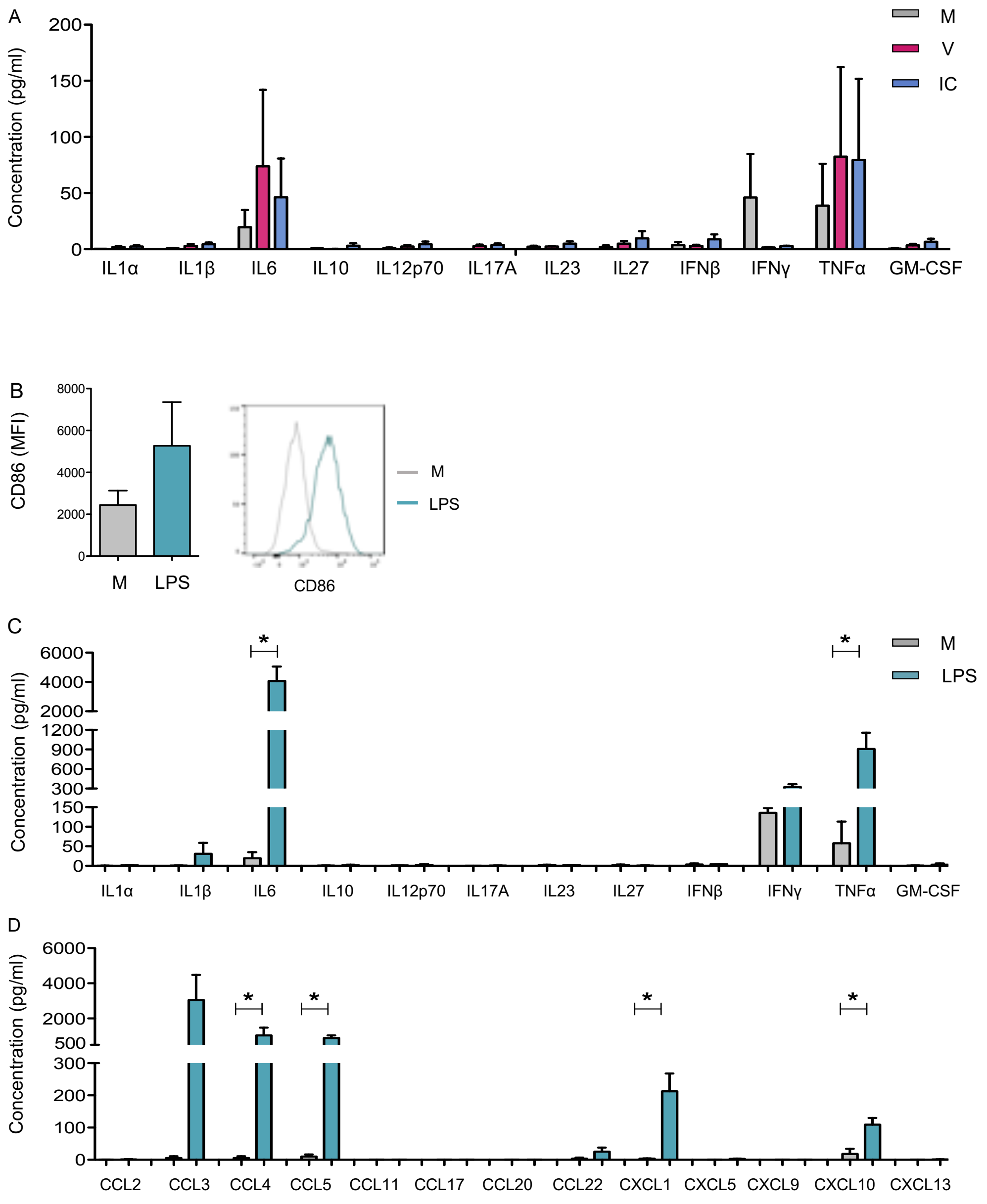


***Supplemental Figure 2:*** **Functional activation of monocytes stimulated with viral determinants or LPS.** BM-derived monocytes were isolated from naive mice and stimulated for 24h *in vitro* with viral determinants (virus or IC) or LPS ($1\mu g/ml$). V, free virions; IC, viral ICs; M, culture medium**. A**. *Cytokine secretion profile of monocytes stimulated by virus (red) o viral ICs (blue) or unstimulated (grey)*. Cytokine release was assessed in supernatants of monocytes isolated from BM of naive mice (>97-98% purity) and stimulated for 24 h. The data represent 6 independent experiments and are expressed as means +/- SEM. Statistical significance was established using a parametric 1-way ANOVA test with Bonferroni's multiple comparisons post-tests (*p < 0.05; **p < 0.01; ***p < 0.001). **B-D**. *Functional activation of monocytes stimulated by LPS*. Activation was assessed by monitoring the CD86 expression level (**B**) as well as the cytokines (**C**) and chemokines (**D**) release monitored in supernatants of monocytes stimulated for 24 h by LPS (green) or left unstimulated (grey). Data represent 5 independent experiments and are expressed as means +/- SEM. Statistical significance was established using a paired Student's t test (*p < 0.05; **p < 0.01; ***p < 0.001).
